# TMEM184b is necessary for IL-31 induced itch

**DOI:** 10.1101/2020.01.25.919902

**Authors:** Erik G. Larsen, Tiffany S. Cho, Matthew L. McBride, Jing Feng, Bhagyashree Manivannan, Cynthia Madura, Nathaniel E. Klein, Elizabeth B. Wright, Hector D. Garcia-Verdugo, Chelsea Jarvis, Rajesh Khanna, Hongzhen Hu, Tally M. Largent-Milnes, Martha R.C. Bhattacharya

## Abstract

Nociceptive and pruriceptive neurons in the dorsal root ganglia (DRG) convey sensations of pain and itch to the spinal cord, respectively. One subtype of mature DRG neurons, comprising about 5% of neurons in the ganglia, is responsible for sensing mediators of acute itch and atopic dermatitis, including the cytokine IL-31. How itch-sensitive (pruriceptive) neurons are specified is unclear. Here we show that *Tmem184b*, a gene with roles in axon degeneration and nerve terminal maintenance, is required for the expression of a large cohort of itch receptors, including those for IL-31, Leukotriene C4, and Histamine. Male and female mice lacking *Tmem184b* show reduced responses to IL-31, but maintain normal responses to pain and mechanical force, indicating a specific behavioral defect in pruriception. Calcium imaging experiments indicate that a reduction in IL-31 induced calcium entry is a likely contributor to this phenotype. We identify an early failure of proper Wnt-dependent transcriptional signatures and signaling components in Tmem184b mutant mice that may explain the improper DRG neuronal subtype specification. Accordingly, lentiviral re-expression of Tmem184b in mutant embryonic neurons restores Wnt signatures. Together, these data demonstrate that Tmem184b promotes adult somatosensation through developmental Wnt signaling and promotion of proper pruriceptive gene expression. Our data illuminate a new key regulatory step in the processes controlling the establishment of diversity in the somatosensory system.

## Introduction

Neurons of the somatosensory system are responsible for detecting temperature, touch, pain, and itch. Somatosensory neurons also relay the first warning signals when peripheral tissues are damaged and local inflammation occurs [7,30,32]. A common consequence of increased skin inflammation is the triggering of itch. Itch-causing pathological conditions have a variety of causes, including skin disorders (atopic dermatitis), drug-induced reactions, systemic disorders such as liver disease, and other neurological disorders [7]. The sensation of itch is distinct from pain and is carried by a subset of itch-specific somatosensory neurons called pruriceptive neurons with cell bodies in the dorsal root ganglia (DRG). Skin itch is triggered by release of pro-inflammatory compounds such as cytokines (IL-4, IL-13, and IL-31), thymic stromal lymphopoietin (TSLP), and histamine from mast cells and epithelial cells, which act upon membrane receptors and channels present in epidermal nerve endings [25,30,32].

Somatosensory neurons display diversity in both protein expression and nerve terminal morphology. They have been classically categorized using soma size, electrical properties (including myelination state), and expression of a few markers that include lectins (IB4), neuroactive peptides, and ion channels. Recently, multiple groups have described the diversity of somatosensory neurons from the adult DRG using single-cell RNA sequencing [20,31]. These studies classified DRG neurons into six to eight groups based on expression of unique transduction or signaling molecules. In addition, the ion channels conferring unique electrical properties have been mapped to eight distinct populations [34]. Finally, developmental single cell profiling of these neurons has enabled a first look at possible developmental trajectories of sensory neurons [28].

One subpopulation of DRG neurons is uniquely identified by expression of the peptides Nppb (natriuretic peptide B) and Somatostatin (Sst), as well as IL-31 receptor type a (Il31ra) [20,31]. These cells, called NP3 or C2 neurons and representing 6-8% of all neurons in the DRG, are purely pruriceptive, transmitting itch, but not pain, signals to the dorsal horn of the spinal cord [12,20,31]. These neurons develop from an initially *Runx1^+^, TrkA^+^* population that ultimately loses *TrkA* expression but gains peptidergic markers such as Calcitonin gene related peptide (CGRP) [17]. How this subset is specified remains a mystery.

Transmembrane protein 184b (Tmem184b) is a member of the Transporter-Opsin-GPCR (TOG) superfamily of proteins [33]. *Tmem184b* participates in axon degeneration following nerve injury, and also maintains proper sensory and motor nerve terminal morphology [2]. In mice lacking *Tmem184b*, nociceptive terminals in the epidermis show swollen endings, and mice show some sensorimotor deficits [2]. The molecular mechanism by which *Tmem184b* exerts its effects is unknown. In this study we sought to identify pathways controlled by *Tmem184b* in neurons. Surprisingly, we found that *Tmem184b* controls the expression of a large cohort of sensory receptors in the DRG, specifically those in the NP3/C2 population mediating pruriception. We found specific behavioral defects in acute itch, but not pain, when *Tmem184b* is absent, showing that *Tmem184b* is required for proper pruriception. *Tmem184b* loss causes a reduction in IL-31-evoked calcium entry in cultured DRG neurons. We show that *Tmem184b* is required in embryonic neurons for expression of critical drivers of sensory neurogenesis, suggesting an early role for *Tmem184b* in sensory development. Restoration of *Tmem184b* in mutant embryonic neurons promotes Wnt pathway gene expression, supporting a role for *Tmem184b* in enabling this essential pathway. *Tmem184b* is also required for proper expression of *Tlx3*, a transcription factor known to promote pruriceptive gene expression. Our data identify a critical role for *Tmem184b* in establishing the proper gene expression repertoire for primary pruriception.

## Materials and Methods

### Animal Models

All animal treatment was approved by the Institutional Animal Care and Use Committee at the University of Arizona (protocol # 17-216). Tmem184b gene-trap mice have been described previously.[2] Mice were bred to *Sst*^Cre^ and Rosa-Flox-stop-Flox-tdTomato (also called Ai9) (lines 013044 and 007909, Jackson Laboratory, Bar Harbor, ME) for histological quantification of the C2 population. Only heterozygous *Sst*^Cre^ mice were used for experiments due to the possible disruption of normal somatostatin expression in homozygous Cre mice.

### RNA sequencing

Total RNA was isolated from adult DRG from 6-month old mutant and wild-type mice (4 per genotype, mixed male and female groups) using the RNaqueous Micro kit (Ambion). All DRGs were pooled for each sample to obtain enough RNA for analysis. Following total RNA extraction, samples were ethanol precipitated to increase purity. Library preparation and sequencing was performed at the Washington University Genome Technology Access Center (GTAC). Data were analyzed using Salmon and DeSeq2 (on our servers or with Galaxy, www.usegalaxy.org). Volcano plots were generated in RStudio; heatmaps of genes for which adjusted P values were less than 0.05 were created using R or with Cluster 3.0 and Java Treeview. To create heatmaps of normalized counts, hierarchical clustering was used in Cluster3.0 to arrange genes by expression similarity. For embryonic DRG analysis, total RNA was isolated using TRIzol and library preparation and sequencing was performed by Novogene; identical analysis methods were used. For bioinformatics analysis, we used Panther’s overrepresentation analysis of biological processes (complete) (http://www.pantherdb.org/) as well as Enrichr (www.enrichr.com). Fold enrichment was calculated as the number of statistically significantly expressed genes from the input dataset divided by the number of genes expected from the entire mouse reference genome. Expected genes in a GO category calculated by applying the proportion of all genes in reference genome associated with a GO category to the number of genes in input dataset. Significance calculated by Fisher’s exact test with a BHM FDR ≤ 0.05. For timecourse and *in vitro* rescue RNAseq analysis, PCA analysis revealed occasional samples that did not cluster with their cohort; these were excluded from downstream DESeq2 analysis while maintaining independent biological replicates of ≥ 3 for all experiments.

To compare *Tlx3* conditional knockout gene expression with that of *Tmem184b*, we obtained raw FASTQ files from NCBI GEO for *Tlx3* mutants (NCBI GEO Accession GSE93394) and ran them through our analysis pipeline using Salmon and DESeq2 prior to comparing datasets.

### Transcription Factor Binding Analysis

Genes identified as differentially expressed by genotype (downregulated in mutant) in E13 ganglia (Adjusted p <0.01) were used to query four databases that predict regulatory element binding by transcription factors. The top 25 hits were chosen from the following four resources: DiRE (top 25 by importance, https://dire.dcode.org/) [9], Enrichr TFs (top 25 by overlap with gene set, https://maayanlab.cloud/Enrichr/) [15], ENCODE (top 25 by overlap with gene set, http://compbio.mit.edu/encode-motifs/) [13], and Chip-Atlas (top 25 by Fold Enrichment in neuronal tissue samples, https://chip-atlas.org/) [26]. If a gene family appeared in 2 or more of these 4 lists, including at least once in the neuronal samples, it was included in the final candidate list.

### Cytokine Injections and Behavior Analysis

Mice were at least 8 weeks old at the time of injection. For itch experiments, both males and females of approximately equal quantities were used and data were pooled. Littermate controls were used, and videos were captured early in the morning to minimize mice falling asleep during videotaping. Mice were acclimated to behavioral chambers (red Rat Retreats, Bioserv) for one hour, removed from the chamber briefly for injections of either IL-31 (Peprotech, 3 nmol in 10 microliters of PBS), chloroquine (200 μg/50 μl), HTMT (0.2 μmol in 25 μl) or 0.9% saline alone, and returned to the chamber, at which time videotaping began. Analysis was done blinded to genotype and injected substance. Scratching bouts were tallied for 30 minutes postinjection. Note that for counting, only scratches directed at the injected area were tallied (neck area) and other non-target directed scratches (for example to the face) were excluded, resulting in numbers that are somewhat lower than those previously reported [3]. If a mouse did not move for 10% or more of the video (≥ 3 min), it was excluded from analysis and assumed to have fallen asleep. Tail flick, hot plate, and Von Frey testing methods have been previously described. [1,4,10] For capsaicin tests, 2 μg of capsaicin in 25 μl, or an equivalent volume of saline, was injected intra-plantarly. Individual mice received either saline or capsaicin on week 1, and then 72 hours later received the other substance into the other hindpaw. Mice were immediately plated in behavioral chambers and watched for 5 minutes by two independent observers to quantify flinches. Counts from each observer were averaged. For itch, pain and mechanical threshold testing, the experimenter was blinded to genotype, and males and females were separated during analysis.

### Motor Function Analysis

Mice were trained on a Rotarod device suspended 40 cm above a padded surface. Rotation speed is 8 rpm. The total time on the rotarod (start time to the time at which the animal falls from the rotarod onto the padded surface) is recorded. Two training sessions are completed, followed by three testing sessions per mouse. Data were averaged per mouse and across genotypes. Male and female performance was evaluated separately, and for each sex, wild type and mutant values were statistically compared using an unpaired t-test in GraphPad Prism.

### Immunohistochemistry and Image Analysis

Isolated ganglia or spinal cords were fixed with 4% paraformaldehyde for 1 hour, immersed in 30% sucrose overnight (4°C), and embedded in OCT cryo-compound (Tissue-Tek^®^ O.C.T.) using isopentane cooled with dry ice. Spinal cords were dissected into segments containing two spinal segments (e.g. cervical 1 and 2). DRG cryosections (14 μm thickness) or spinal cord sections (20 μm thickness) were-mounted onto charged microscope slides. Following washes in PBST and 1 hour of blocking in 5% goat serum in PBST, sections were incubated overnight at 4°C with rabbit NeuN (1:250, Proteintech), or for 1 hour with Alexa Fluor^®^ 488 Mouse anti-β-Tubulin, Class III (BD Pharmingen). Secondary antibody for NeuN was goat anti-rabbit Alexa Fluor 633 (Thermo Fisher). Sections were mounted in Vectashield (Vector Biolabs). Spinal cords were imaged with a ZEISS AxioZoom V16 Fluorescent Microscope. DRGs were imaged using a ZEISS Axio Observer Z1 or LSM880 inverted confocal microscope. For DRG counting, only neurons with a visible NeuN-positive nucleus in the section were counted. For each genotype, n = 5 mice. For each mouse, ≥800 neurons were counted from at least 3 separate ganglia and at least 3 sections per ganglia. Due to the small neuron numbers in some ganglia, we occasionally used up to 25 individual sections from up to 6 ganglia from a single mouse.

### LacZ staining

Adult mice were euthanized with CO2. Hairy skin from the top of each foot was removed and fixed in 100% acetone for 8 hours at room temperature, followed by immersion in 1mg/ml X-gal in staining solution (0.1 M Phosphate Buffer pH 7.5, 5 mM potassium ferricyanide, 5 mM potassium ferrocyanide, 20 mM Tris-HCL, 0.02% NP-40, 0.01% sodium deoxycholate) for 48 hours at 4°C. Tissues were then rinsed, post-fixed in 4% paraformaldehyde/1xPBS overnight, washed in PBS, submerged in 30% sucrose/1xPBS, and embedded in O.C.T. media. Sections of 15-20 μm thickness were cut and collected on glass slides and kept frozen until use.

Antibodies on these sections included Rat anti-CD3 epsilon (1:100, NBP1-26685, Novus Bio) and Rabbit anti-C-Kit (1:100, D13A2, Cell Signaling Tech).

### Neuronal Culture

For adult neurons, ganglia from all spinal levels were pooled in DMEM on ice, followed by digestion with Liberase TM (Roche) and 0.05% Trypsin (Gibco). Neurons were dissociated with a P1000 pipette tip, spotted in dense culture spots (20uL) on poly-D-lysine (Sigma) and laminin coated chambered coverglass (Nunc) or 100mm glass coverslips, and grown overnight in Dulbecco’s Modified Eagle Medium (Gibco) with 10% fetal bovine serum (Atlas Biologicals) and Penicillin/Streptomycin (Gibco). For embryonic DRG culture, ganglia were dissociated with Trypsin, triturated with a P1000 pipette tip, and plated in 5-10 μl spots in a 24-well dish previously coated with poly-D-lysine and laminin. Media contained B27 (Gibco), 5-fluoro-deoxy-uridine (FDU) and nerve growth factor (NGF) (Invitrogen). Half of the media volume was exchanged every 5 days until cells were collected for analysis.

### Calcium Imaging

Adult DRG neurons were imaged either with Fluo-4 dye on a Zeiss Observer Z1 microscope, or with Fura-2 dye on an Olympus BXW microscope under a 10X immersion objective lens with a filter wheel and Hamamatsu camera, each with a frame rate of one image/3 sec. Data shown used ratiometric, perfusion-based Fura-2 imaging with the exception of IL-31 and chloroquine, which used Fluo-4 due to quantity constraints (IL-31) or autofluorescence in UV wavelengths (chloroquine). Fluorescence videos were acquired via Zeiss or HCImage Software, processed and analyzed using MATLAB, ImageJ/Fiji, and RStudio for quantification of fluorescent responses. We used a custom-written R script to identify neurons via responses to high potassium in each experiment. Responses to individual agonists were determined manually for all neurons using the following criteria: peak response must change by 10% from baseline, must initially show calcium rise in the agonist application time window, must increase with an upward deflection in slope of at least 45 degrees rather than a gradual baseline shift, and display a waveform similar to that previously reported. Each agonist was evaluated using cultures from at least 3 mice, and all experiments contained greater than 100 neurons analyzed. Coverslips with extensive motion or other artifacts were excluded from analysis. Perfusion artifacts (single frame spikes) were identified and removed using low-pass filtering in R prior to analysis.

### Luciferase Assays

HEK293T cells (ATCC) were transfected with M50TOP-Flash and plasmids expressing either Wnt3a (Addgene) or human TMEM184b with a Myc-tag. Both plasmids contained a fluorescent marker (GFP or Venus, respectively) to assay transfection levels. Cells were split to poly-D-lysine coated 96-well white plates at 24 hours post-transfection, and assay was conducted 48 hours post-transfection using a plate reader (BMG FLUOstar). Luciferase activity was measured by lytic assay using NanoLuc (Promega Nano-Glo kit). Viafluor 405 was used to normalize for number of cells per well. pAd-Wnt3a was a gift from Tong-Chuan He (Addgene plasmid # 12518).

### Experimental Design and Statistical Analysis

All statistical analyses were performed in GraphPad Prism except for weighted linear regression analyses, which were performed in R. Statistical methods specific to individual experiments are included in figure legends or descriptions of individual experiments in the Methods section. Where applicable for multiple test corrections, all False-Discovery Rates determined with Benjamini-Hochberg Method (BHM); thresholds set at (FDR) ≤ 0.05. For all figures, asterisks indicate p < 0.05 (*), p < 0.01 (**), p < 0.001 (***), or p < 0.0001 (****).

Estimating proportions of neurons responsive to various agonists depends on the highly variable sampling of neurons from each mouse. Thus, weighted linear regression was employed to estimate means of population percentages between genotypes. Weights, *w*, for each mouse of both genotypes per agonist, *i*, consisted of the number of neurons, *n*, sampled from a given mouse, *j*, divided by the sum of all of the neurons exposed to a given agonist. These weights were scaled by 10 to provide an “effective mouse weight.”

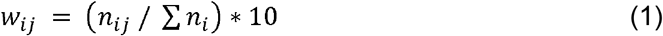

The proportions of responding neurons were then regressed to estimate the means between genotypes. Magnitudes were quantified relative to a baseline fluorescence. Because of the sampling variability of responding neurons between preparations, these magnitudes were weighted as above in Eq. 1. However, mouse weights depended on the number of responsive neurons, as the strength of responding neurons were the measurements being evaluated.

### Data accessibility

The data sets generated and analyzed in the current study have been deposited to NCBI GEO at the following links: GSE154435, GSE154354, GSE154316.

### Code accessibility

All custom-written code in R and MATLAB for calcium imaging and RNAseq analysis is available at GitHub (https://github.com/eriklarsen4/ggplot-scripts.git and https://github.com/martharcb/Fluo4_R).

## Results

### Tmem184b is Required for Expression of Pruriceptor-Specific Markers

In studying the nervous system phenotypes of loss of the transmembrane protein *Tmem184b*, which include axon protection as well as dysmorphic nerve terminals [2], we sought to identify expression changes that could contribute to these phenotypes. We performed RNA sequencing on isolated adult DRGs of 6-month-old *Tmem184b* gene trap mutant mice, in which Tmem184b transcripts are globally reduced by 95% [2], and compared transcript abundance to age-matched wild-type mice. A total of 406 transcripts were differentially expressed (Fig. 1A and Table, Supplemental Digital Content 1). Strikingly, the downregulated genes contained a large fraction of markers previously identified in single-cell RNAseq studies as being unique to a population of pruriceptive neurons (called NP3 or C2 in previous studies) [5,21,31]. The key markers of NP3/C2 neurons are dramatically downregulated, with reductions in *Il31ra, Nppb*, and *Sst* being among the largest changes seen in the dataset (Fig. 1B). In addition to NP3/C2 genes, other types of neurons also showed decreased expression of their unique markers, including NP2/C4 (identified by *Mrgpra3*) and NP1/C5-6 (identified by *Mrgprd*) (Fig. 1C). Taken together, this data identifies *Tmem184b* as a major regulator of somatosensory gene expression, particularly in pruriceptive populations.

**Figure 1.**
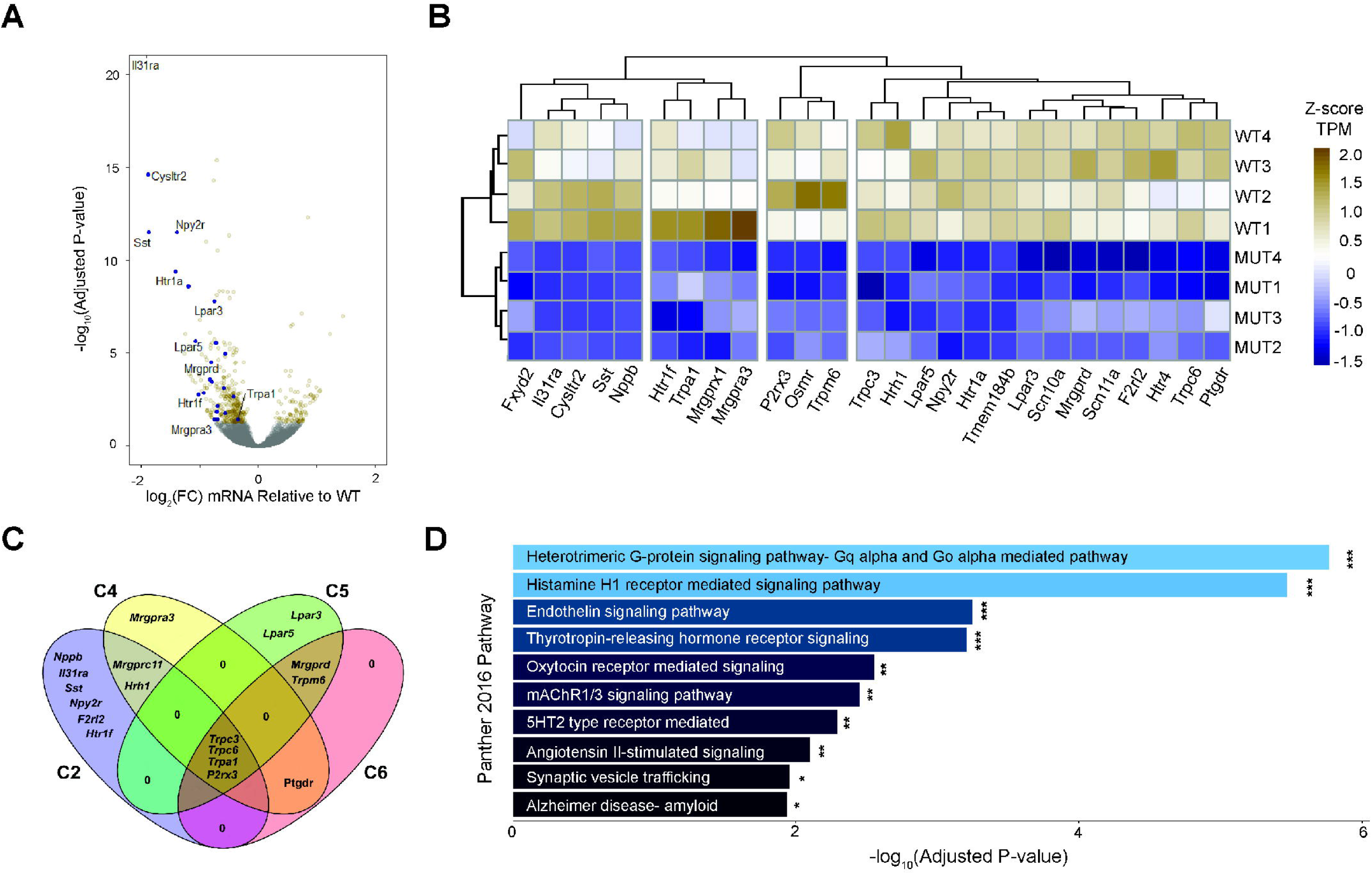
Tmem184b is Required for Expression of Pruriceptor-Specific Markers. (A) Volcano plot of RNA sequencing data. Gold, differentially expressed (DEGs) (n = 381 genes); navy shows a subset of itch-related DEGs (n = 24 genes); gray, non-differentially expressed genes (n = 12,550 genes). Benjamini-Hochberg Method-derived FDR; FDR ≤ 0.05. (B) Heatmap showing aligned, variance-normalized samples (N = 4 per genotype), and DEGs of interest independently clustered by expression similarity. (C) Venn diagram of selected sensory neuron expressed DEGs showing their specific expression in subsets of nociceptors. (D) Genetic labeling of Sst-Cre somatosensory neurons (magenta) expressing tdTomato in Tmem184b gene trap mice. Green indicates NeuN (neurons). (E) Percent of neurons labeled with Sst-Cre>Ai9 in wild type and Tmem184b-mutant mice. N=5 mice and ≥800 neurons analyzed per mouse. Data presented as mean ± SD (7.20 ± 2.00, 4.21, ± 1.74, t = 2.52, p = 0.036, unpaired t test) (F) Over-represented Panther pathways (FDR ≤ 0.05) in downregulated DEGs.

To predict the functional significance of these expression changes, we performed pathway analysis using Panther. Downregulated transcripts were enriched in pathways such as GPCR signaling, serotonin receptor signaling, and synaptic transmission (Fig. 1D). When human phenotype ontology was examined, pruritus was the top and most significantly changed feature (p < 0.003). These data show that *Tmem184b* is required for appropriate transcription of signaling pathways critical to normal sensory function.

### *Tmem184b* is required for IL-31 induced itch, but not pain, responses

To test the behavioral consequences of the expression changes observed in *Tmem184b-mutant* mice, we challenged them with agonists that promote itch or pain. The cytokine IL-31, implicated in atopic dermatitis, activates the IL31RA/OSMR heterodimeric receptor on NP3/C2 DRG neurons, whereas the anti-malarial agent chloroquine (CQ) activates MRGPRA3 on NP2/C4 DRG neurons [3,6,22]. *Tmem184b*-mutant mice show a reduction in target area scratching evoked by IL-31 injection compared to wild type mice (Fig. 2A). Mutant mice also show a small but not statistically significant reduction in scratching to the H1-Histamine receptor agonist HTMT (Fig 2B) but no apparent deficit in their response to the MRGPRA3 agonist chloroquine (Fig. 2C). We do not see a deficit in pain responses to intraplantar injection of the TRPV1 agonist, capsaicin (Fig. 2D), but observed that mutant male mice responded more strongly to capsaicin than wild type males. To examine more broadly the effects on nociception and mechanoreception, we performed thermal pain and mechanical threshold testing. We also evaluated the motor coordination of mutant mice. *Tmem184b*-mutant mice responded normally in all of these tests (Fig. 2E-H). Taken together, this data indicates that *Tmem184b* preferentially impacts behavioral responses to IL-31.

**Figure 2.**
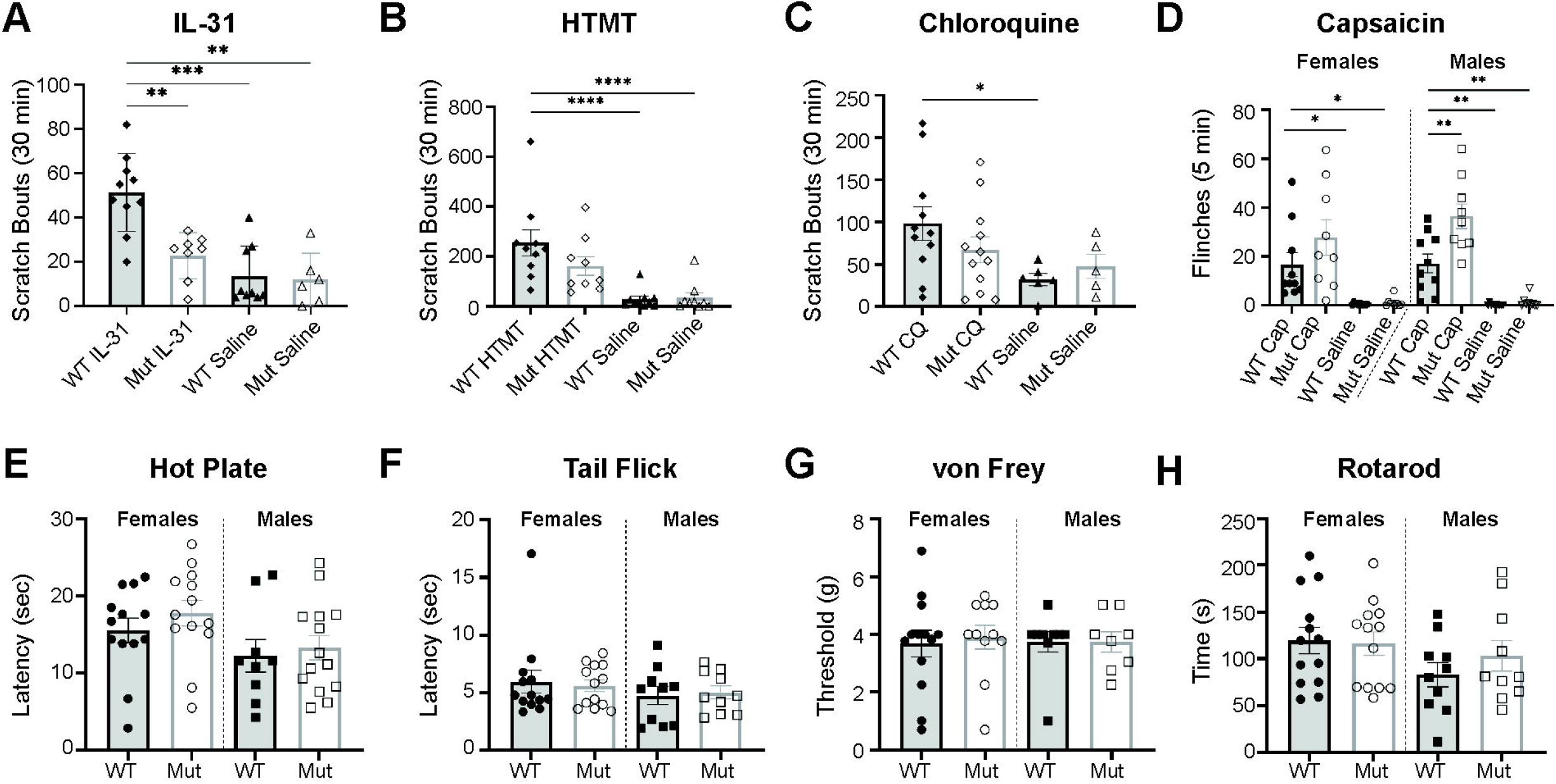
Tmem184b is required for itch, but not pain, responses. Wild type measurements use filled bars and markers; mutant measurements use open bars and markers. For (A-C), n= 8-12 for itch agonists and 5-8 for saline controls. Significance evaluated using two-way ANOVA with Dunnett’s post-hoc test. Symbols: ***(p<0.001), **(p<0.01), *(p<0.05). (A) IL-31 induced scratching (p = 0.0096 between WT IL-31 and mutant IL-31, 2-way ANOVA with Dunnett’s post-hoc test). (B) Histamine trimethyl toluidide (HTMT)-induced scratching (p = 0.098 between Wt and mutant HTMT-injected mice). (C) Chloroquine-induced scratching responses (p = 0.31 between WT and mutant CQ-injected mice). (D) Capsaicin intraplantar injection and flinching counts, separated by sex (female cap comparison, p = 0.23; male cap comparison, p = 0.0014). Each sex separately evaluated using 2-way ANOVA with Dunnett’s post-hoc test. N=9-10 animals per sex and genotype analyzed. (E-H), n=8-13 mice per sex and genotype. All data presented as mean ± SEM. Analysis by two-tailed, unpaired t-test within sex. (E) Hot plate analysis (female, p = 0.3404; male, p = 0.694). (F) Tail flick analysis (Female: p = 0.762; Male: p = 0.769). (G) Baseline mechanical thresholds (Female: p = 0.735; Male: p = 0.983). (H) Time spent on rotarod (Female: p=0.87; Male: p=0.35).

### Tmem184b is expressed in hair follicles, but not T or mast cells, in hairy skin

Our RNAseq was performed on sensory ganglia, far removed from the site of pruriception in the skin. We wanted to assess the likelihood that other cells (beyond neurons) could be affected in our mutants and contribute to the reduction in IL-31-induced scratching. IL-31 is produced primarily by Th2 cells and can activate IL31RA/OSMR dimeric receptors on mast cells and keratinocytes, in addition to sensory neurons, in the skin [24]. Mast cells in the skin can also activate the NP3/C2 population through release of leukotrienes, serotonin, and other agonists [30]. To begin to answer this question, we asked whether *Tmem184b* was expressed in pruritus-relevant cell types in skin. We took advantage of the lacZ reporter in our gene-trap mutant, which is expressed in the pattern of endogenous *Tmem184b*. In hairy skin, TMEM184B is expressed in hair follicles (Fig. 3A) but not T cells (CD3+) or mast cells (c-Kit+) (Fig. 3B-D). Antibodies made to Tmem184b from commercial vendors (Sigma), as well as two separate attempts at custom-made antibodies, still detect bands in gene trap mice by Western blot (data not shown), indicating non-specificity of binding. Therefore, we are unfortunately unable to see the localization of endogenous protein in the skin or at terminals. However, our previous work using lacZ staining shows that TMEM184b is expressed in most, if not all, neurons in the mouse DRG [2]. Therefore, while it is possible that TMEM184b has non-cell autonomous effects in the skin, we suspect that our phenotypes result primarily from sensory neuron reduction of IL31RA/OSMR signaling.

**Figure 3.**
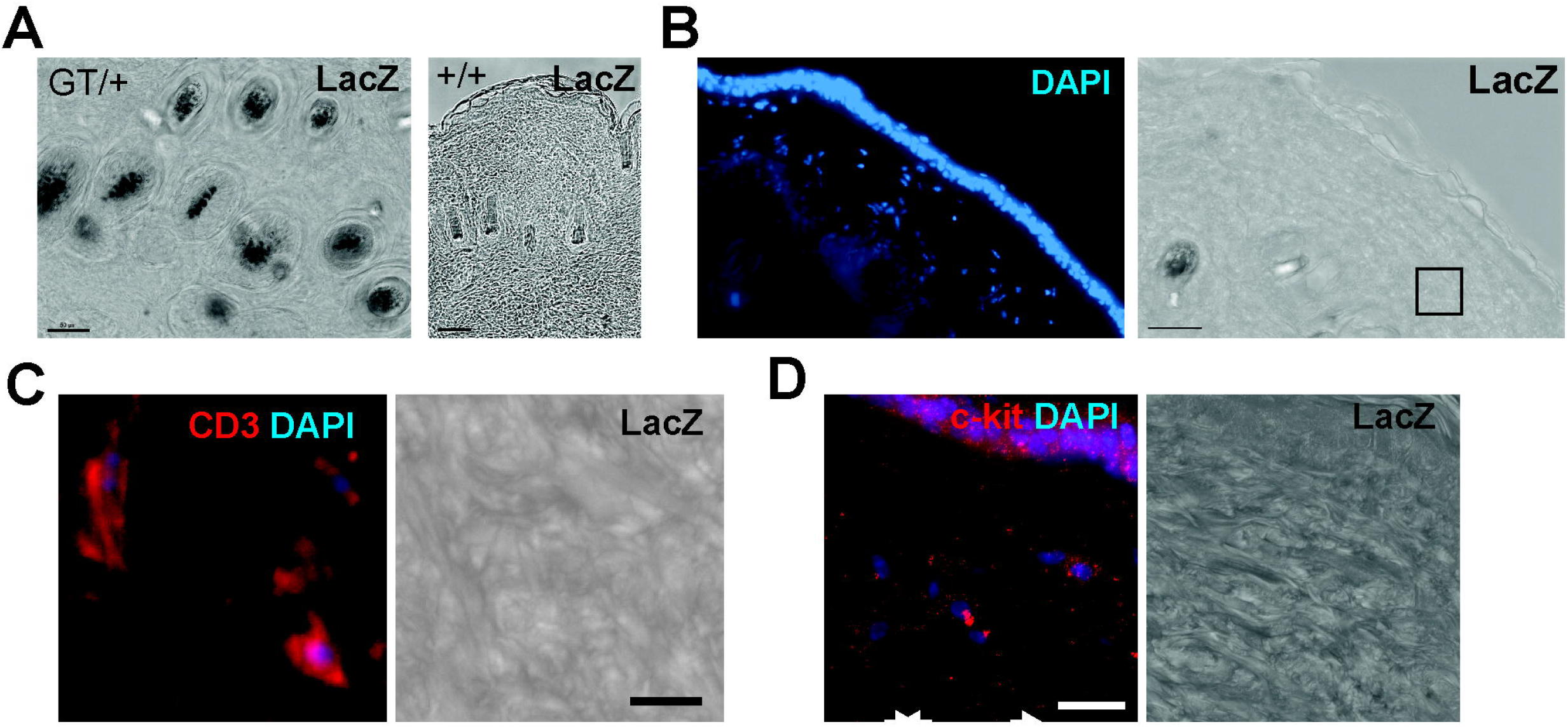
Tmem184b is expressed in hair follicles, but not T or mast cells, in hairy skin. All panels show representative images from heterozygous mice (GT/+) except for the right panel of part A (+/+). The gene trap allele expresses β-galactosidase under the control of the Tmem184b promoter, allowing labeling of cells expressing Tmem184b. A, X-gal reactivity (dark) showing hair follicle expression of Tmem184b in GT/+ mice. No background X-gal reactivity is detected in hairy skin of +/+ mice. Scale bars = 50 μm. B, cross-section of hairy skin showing both the dermis and epidermis, illustrating the lack of X-gal reactivity in all locations aside from hair follicles. Scale bar = 50 μm. C, immunohistochemistry with CD3, a T-cell marker, showing the locations of T-cells in the dermis and their lack of X-gal reactivity. Pictures are from the boxed area in B. Scale bar = 10 μm. D, immunohistochemistry to c-Kit, a marker of mast cells, showing lack of X-gal reactivity. Note that c-kit also labels melanocytes (upper right of the images). Scale bar = 20 μm.

### IL-31 evokes less calcium entry in *Tmem184b*-mutant DRG neurons

To determine whether NP3/C2 neurons are still present in Tmem184b mutant mice, and to evaluate any difference in the quality of their responses to NP3/C2 agonists, we performed calcium imaging on dissociated adult DRG neurons (Fig. 4 and Table, Supplementary Digital Content 2). In wild type neurons, IL-31 evokes relatively small calcium transients (Fig. 4A-B), consistent with previous reports [3]. Because DRG responses are graded, we considered whether changes in amplitude or in total calcium entry could explain the decrease in IL-31-evoked scratching. To measure response magnitudes, we analyzed responses following loading with Fura-2 ratiometric calcium dye, as it is sensitive to small changes in calcium. Among neurons classified as responders, IL-31 responders showed a trend toward lower peaks (Fig. 4B-C), and their overall calcium entry for the duration of application was significantly reduced (Fig. 4D). The reduction in calcium entry of individual neurons to IL-31 thus may contribute to the decrease in IL-31-induced pruriception in adults. Interestingly, in isolated adult ganglia IL31RA trends toward only a slight decrease at the protein level (Fig. 4E). However, in considering the mechanisms of IL-31RA signaling, we identified its co-receptor OSMR, as well as Jak1 and Stat3 in our differentially expressed genes (all are downregulated) (Table 1, Supplementary Digital Content 1). Our data is consistent with reduction in expression of many genes in the IL-31RA/OSMR signaling pathway in Tmem184b mutant mice, that together lead to a behavioral deficiency in IL-31 evoked itch.

**Figure 4.**
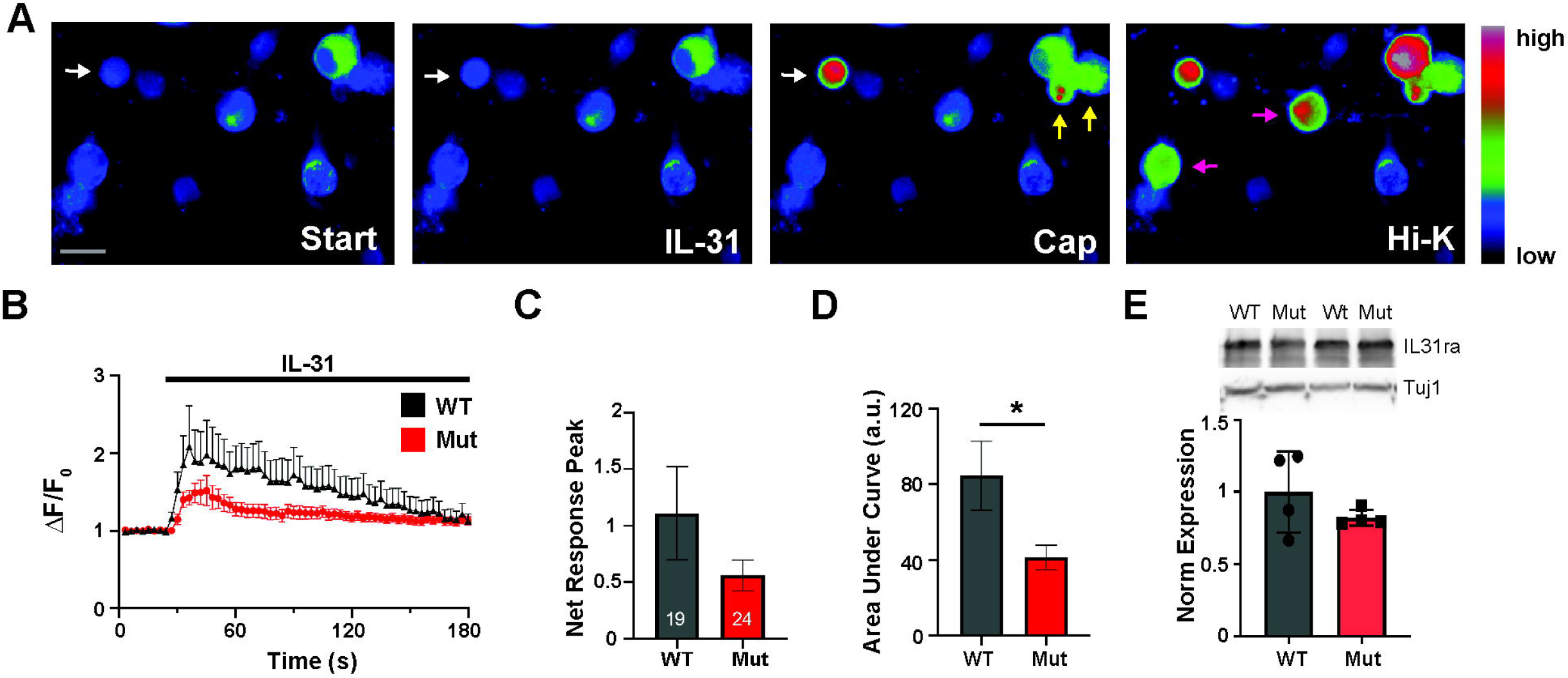
IL-31 evokes less calcium entry in Tmem184b mutant DRG neurons. (A) Representative images from a wild type mouse showing neurons responding to IL-31 (1μM), capsaicin (1μM), and/or High-K+ (70 mM KCl) Ringers solution. Scale bar, 20 μm. (B) Average IL-31 response calculated from wild type (black) and mutant (red) neurons classified as IL-31 responders. Wild type (n = 19) and mutant (n = 24) responding neurons cultured and imaged on the same day with the same solutions were averaged. (C) Net responses (where pre-stimulus baseline is set to 0) of cells from B. Error bars show SEM; p = 0.17, unpaired t-test. (D) Area under curve analysis for IL-31 stimulus window of cells from B. Error bars show SEM; p = 0.02, unpaired t-test. Experiments used either Fluo-4 AM dye (A) or Fura-2 AM dye (B-D). (E) Western blot showing IL31ra levels in adult ganglia from wild type (WT) and Tmem184b gene trap mice (Mut). Beta-III tubulin (Tuj1) was used as a loading control. N = 4 mice per genotype are included in the quantification.

We also quantified percent responders and peak responses to NP3/C2 agonists CYM5442 (an S1PR1 agonist) and LY344864 (an HTR1F agonist); NP2/C4 agonist, chloroquine (an MRGPRA3 agonist); C5/C6 agonist, beta-alanine (an MRGPRD agonist); and broad nociceptive agonists, capsaicin (a TRPV1 agonist) and allyl isothiocyanate (AITC, a TRPA1 agonist). We did not observe significant shifts in the percentage responders of any neurons in these categories (Table, Supplementary Digital Content 2). Furthermore, their response peaks did not differ from wild type mice. These results indicates that most pruriceptive neurons are still present and functional in Tmem184b mutant mice. When considering that all of the above receptors except IL31RA are reduced by less than 50% at the transcript level by RNAseq (Table, Supplementary Digital Content 1), these results are not surprising and indicate that this level of change is not sufficient to compromise cellular sensitivity to these agonists.

### *Tmem184b* regulates key developmental pathways in somatosensory neurons

We wanted to take a broader look at how TMEM184b contributes to the function of sensory ganglia. We first asked if there were TMEM184b-dependent alterations in gene expression during embryonic sensory neuron development. In mice, unique DRG subpopulations arise from a uniform group of neural crest precursors over a period of about three weeks in late embryonic and early postnatal life (from E10-P10) [17]. *Tmem184b* is expressed in DRG neurons as early as E11.5 according to a recent single-cell RNAseq study [28]; we independently confirmed its expression at E13 in somatosensory ganglia (Fig. 5A and Table, Supplementary Digital Content 3). To identify developmental pathways dependent on *Tmem184b*, we isolated RNA from embryonic day 13 (E13) whole dorsal root ganglia and compared transcriptional signatures between mutant and wild type ganglia using RNAseq. We identified 1635 genes for which transcripts significantly changed in the absence of *Tmem184b* at this timepoint (Adjusted P < 0.01) (Fig. 5A). Using pathway analysis, we identified multiple key developmental processes, including axonogenesis and neuronal development and differentiation, that are substantially affected by the loss of *Tmem184b* at E13 (Fig. 5B-C). These results underscore that *Tmem184b* controls key aspects of the establishment of the somatosensory system.

**Figure 5.**
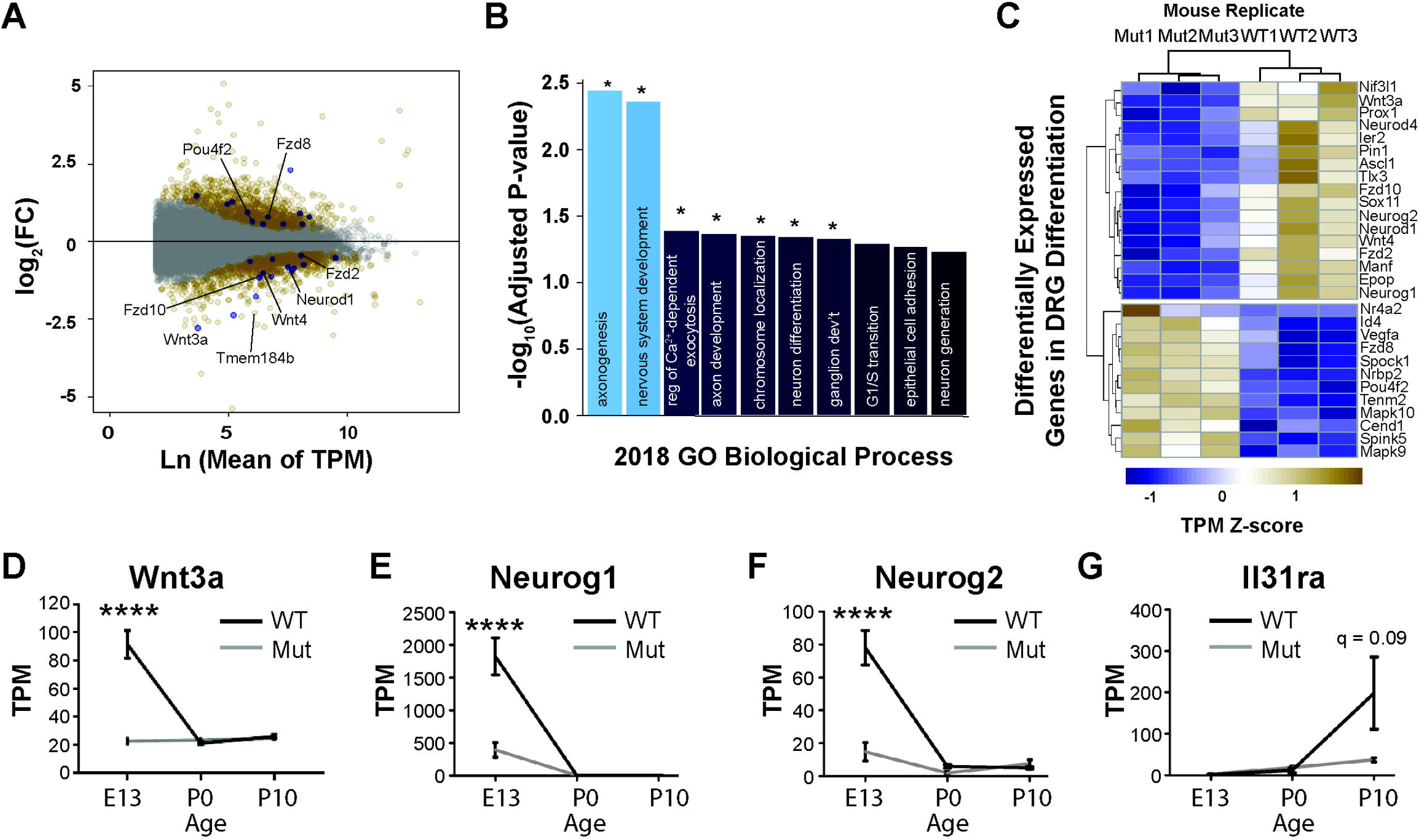
*Tmem184b* regulates key developmental pathways in somatosensory neurons. (A) MA plot of RNAseq E13 *Tmem184b* GT/GT dorsal root ganglia, showing fold change versus average transcript expression across all replicates. Colors: gold, differentially expressed genes (DEGs); blue, neuron differentiation DEGs as annotated in gene ontology (Panther); gray, non-DEGs. (B) Enriched GO Biological Processes in E13 differentially expressed genes (gold from D). Top 10 most significantly enriched pathways (by adjusted P-value) are shown. *, FDR < 0.05. (C) Heatmap of selected, differentially expressed neuron differentiation genes at E13. (D-G) Normalized counts of individual genes from RNAseq analysis of E13, P0, and P10 mutant and wild type mouse dorsal root ganglia. n = 3 mice per time-point and genotype. Asterisks indicate statistical significance as calculated by DESeq2 (Wald Test with FDR ≤ 0.05). Data presented as mean ± SEM. (D) *Wnt3a* (q < 0.0001). (E) *Neurog1* (q < 0.0001). (F) *Neurog2* (q <0.0001). (G) *Il31ra* (q =0.09).

One of the main pathways controlling neurogenesis and differentiation of DRG is Wnt signaling. Neuronal precursors undergo two waves of neurogenesis stimulated by Wnt pathway activation [8,18]; the later wave produces the majority of nociceptors and pruriceptors. Wnt signaling promotes the expression of transcription factors critical in establishing neuronal numbers and identity in migrating neural crest progenitors, including Neurog1 (*Ngn1*) and Neurog2 (*Ngn2*) [14,17,18,27]. In *Tmem184b-mutant* ganglia, we observe striking reductions at E13 in transcripts for *Wnt3a, Ngn1*, and *Ngn2* (Fig. 5C-F), suggesting that an initial failure in Wnt expression could produce deficiencies in the transcriptional programs necessary for nociceptor and pruriceptor development. *Tmem184b* is required for the proper transcript expression of components at all levels of the Wnt pathway, including ligands (*Wnt3a, Wnt4*), receptors (*Fzd2, Fzd10*), and transcription factors (*Tlx3, Neurod1*) (Fig. 5C and Table, Supplementary Digital Content 3).

To analyze the trajectories of somatosensory neuron populations as they are adopting their adult fates, we combined our E13 analysis with a parallel transcriptomic evaluation of gene expression in developing ganglia at postnatal day 0 and postnatal day 10. At the latest time point, expression of NP3/C2 DRG markers has been previously observed [17]. Because Wnt3a, Neurog1 and Neurog2 primarily express during embryonic stages, they are only affected at E13 (Fig. 5D-F). In contrast, *Il31ra* transcript expression initiates between birth and P10 in wild type but remains at very low levels in *Tmem184b* mutants (Fig. 5G). Many genes necessary for somatosensory development or mature function are altered at one or more timepoints during this critical developmental window (Table, Supplementary Digital Content 3). Taken together, our data support a model in which *Tmem184b*, acting early in neuronal development, controls a Wnt-dependent developmental program leading to proper gene expression and function of pruriceptive neurons.

### *Tmem184b* controls expression of Wnt signaling components in embryonic DRG neurons

To more directly test if TMEM184B activity in DRG neurons controls critical developmental gene expression networks, we took advantage of the simplicity of embryonic DRG cultures to evaluate the effects of re-expressing *Tmem184b* in mutant neurons. We cultured E13 embryonic DRG neurons from wild type or mutant mice for 14 days (DIV 14, roughly equivalent timewise to postnatal day 8) (Fig. 6A). In a subset of neurons from each mutant embryo, we re-expressed *Tmem184b* using lentiviral infection on the day of dissociation (Fig. 6A-B). At day 14, we collected RNA and performed RNA sequencing for each condition. Expression of NP3 neuron marker genes did not occur normally in this *in vitro* system after 14 days (Fig. 6C), therefore it is likely that non-neuron-intrinsic factors influence the final differentiation of this population, a finding in line with previous work [28]. However, we were still able to directly test the effects of restoration of *Tmem184b* on overall developmental gene expression. We identified differentially expressed genes in mutant neurons relative to wild type, and in mutant neurons with and without restoration of *Tmem184b* (Fig. 6D-E). Within these genes, we identified those that both decreased in the absence of *Tmem184b* (Fig. 6D, blue dots) and increased in mutants upon re-expression of *Tmem184b* (Fig. 6E, gold dots). Our analysis identified 304 genes matching this pattern. Gene ontology analysis revealed biological processes significantly over-represented by these genes, representing processes positively regulated by TMEM184B activity (Fig. 6F). The most over-represented process among genes positively regulated by *Tmem184b* is the planar cell polarity (PCP) pathway (23.5-fold enriched). Canonical Wnt signaling was also enriched in this group of genes (7.93-fold). *Tmem184b* regulated expression of components at multiple levels of the Wnt pathway, including receptors (*Fzd1, Fzd10*) as well as transcription factors (*Tcf7l2, Taz*) (Fig. 6G), matching our results from whole ganglia at E13 (Fig. 5). Many more genes that positively influence Wnt signaling (green) were identified as compared to those that inhibit Wnt signaling (red). Of note, because these cultures were significantly enriched in neurons over glia (due to the presence of the anti-mitotic factor FdU for the duration of culture), it suggests that the expression changes to Wnt components are likely occurring within the DRG neurons themselves. Many of the processes controlled by *Tmem184b* also influence critical neuronal functions including calcium regulation (7.86-fold enrichment) and axon guidance (4.18-fold enrichment). This data implicates *Tmem184b* in controlling key developmental steps in the establishment of the somatosensory system.

**Figure 6.**
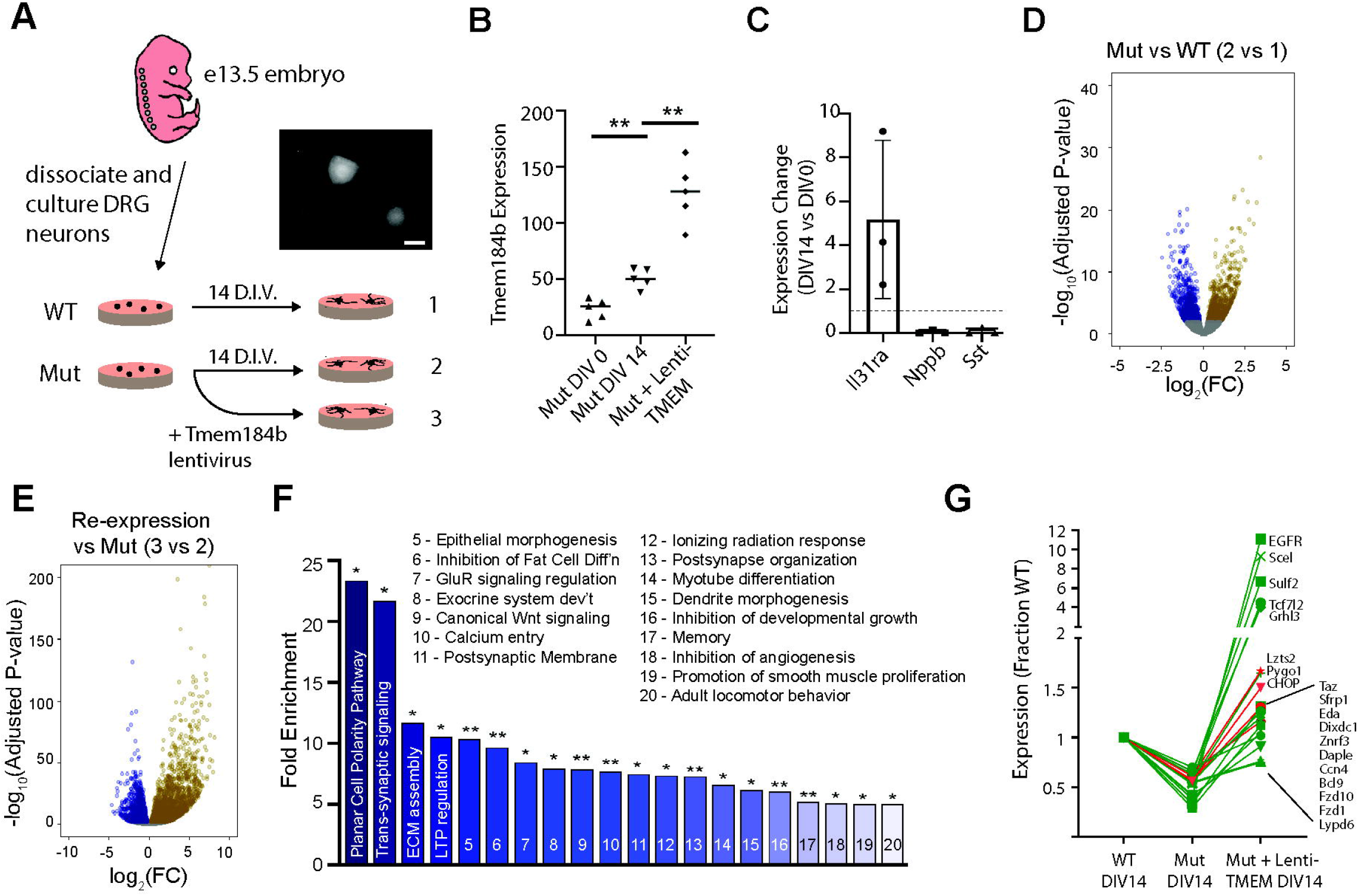
Tmem184b controls expression of Wnt signaling components in embryonic DRG neurons. (A) Schematic of the experimental design. Samples from dissociated DRG neurons at DIV 0 (N = 4 per group) and DIV 14 (N = 5 per group) were collected for RNAseq analysis. Inset, lentivirus infection of mutant eDRG neurons as indicated by IRES-Venus expression at DIV14. At this endpoint, around 50% of neurons showed Venus expression. (B) Evaluation of *Tmem184b* expression level in rescue experiments through analysis of mean normalized counts across samples at DIV14. (C) qRT-PCR analysis of the indicated genes in wild type embryonic DRG cultures at DIV 14 vs DIV 0 (dissociated cells on the day of culture). (D) Volcano plots showing (left) DIV14 Tmem184b-mutant vs DIV14 WT and (right) DIV14 rescue vs DIV14 mutant transcript expression. Gray dots show genes not significantly altered. Navy, downregulated genes (Adj P ≤ 0.01); gold, upregulated genes (Adj P ≤ 0.01). (E) Genes both downregulated in DIV14 mutant eDRGs (B, navy) and upregulated in rescued DIV14 mutant eDRGs (C, gold) were analyzed using Panther gene ontology over-representation analysis. Fold enrichment when compared to all mouse transcripts is shown for all processes with enrichment scores >5. *, FDR < 0.05 and **, FDR <0.01. (F) Expression of genes annotated as Wnt signaling effectors, normalized to WT DIV14 levels. Green genes positive regulate Wnt signaling (15 genes), while red genes negatively regulate Wnt signaling (4 genes).

### Effects of Tmem184b on Wnt Pathway Activation are Indirect

To better understand how Tmem184b exerts these early transcriptional effects, we identified enriched transcription factor binding sequences in downregulated genes from E13 ganglia. To meet our criteria for likely candidates, transcription factors had to appear in the top 25 results of at least 2 of 4 different analyses (Enrichr, ENCODE, DIRE, Chip-Atlas), have been identified in Chip-Seq analyses from neuronal tissues, and have significant p-values for enrichment (p<0.05). Using these strict criteria, we identified four families of transcription factors that are likely targets of Tmem184b downstream signaling (Fig. 7A). Remarkably, two of these are known downstream mediators of Wnt signaling, including both the canonical and Planar Cell Polarity pathways (TCF and FOS/JNK, respectively). The others, NFY and MYC, are involved in growth and metabolism, critical processes during the expansion of neuronal progenitors. This analysis supports a role for Tmem184b in positively regulating Wnt signaling and early developmental decisions.

**Figure 7.**
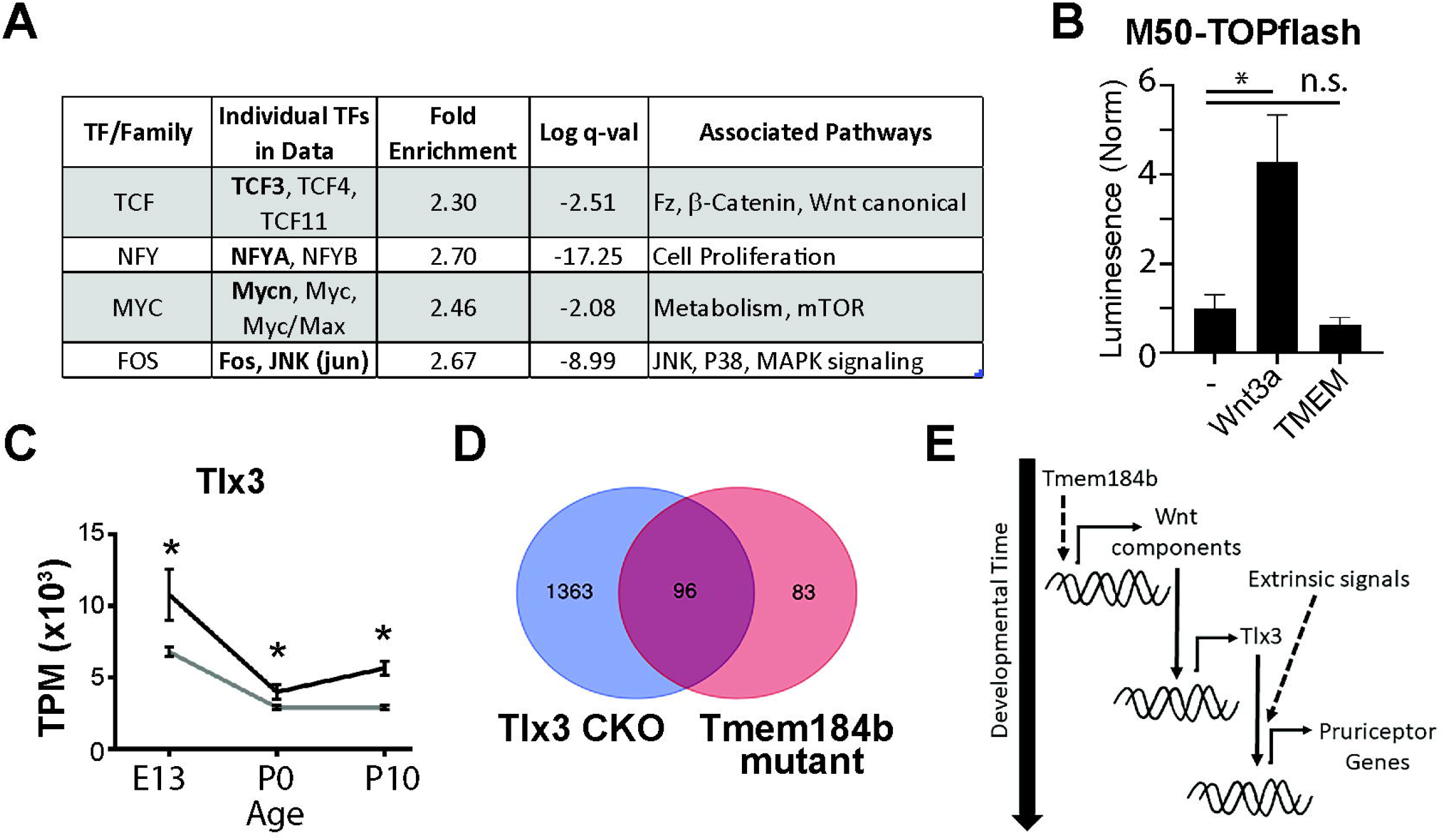
Tlx3 is a candidate linking embryonic gene expression to proper pruriceptor differentiation. (A) Candidate transcription factor (TF) families enriched at E13 differentially expressed gene promoters. Bold gene names in individual TFs column indicate presence in publicly available Chip-Seq data from neuronal tissues. Fold enrichment and log q-value are the maximum and minimum values, respectively, for TFs found in neuronal Chip-Seq experiments. (B) Luciferase assay in HEK293T cells transiently transfected with M50TOP-Flash (canonical Wnt reporter) and the indicated expression constructs, n = 3 independent transfections. p = 0.038 for control vs Wnt3a; p = 0.35 for control vs TMEM184b (student’s t-test). (C) Normalized counts for Tlx3 (q < 0.05 for all timepoints). (D) Venn diagram showing overlapping differentially expressed gene count between Tlx3F/F, NaV1.8-Cre conditional knockout adult DRG (blue) [11] and Tmem184b gene trap mice (red, adjusted p<0.01). (E) Proposed model of Tmem184b-dependent pathways showing the sequential activation of transcription factors that enable pruriceptive gene expression.

We next asked if Tmem184b can directly activate transcription downstream of Wnt signaling. Using an established luciferase assay reporter for canonical Wnt signaling, we observed increased promoter activity when HEK293T cells were transfected with Wnt3a, but not TMEM184b (Fig. 7B). Therefore, while components of the canonical Wnt signaling pathway are expressed and can be activated in this heterologous system, over-expression of Tmem184b is unable to do so directly.

What is the relationship between Tmem184b’s control of early and late transcriptional events? Very few individual differentially expressed genes overlap between E13 and adult data sets, suggesting that a developmental relay mechanism could be at play. A clue to this mechanism comes from examination of the expression of Tlx3, a homeobox transcription factor known to promote the specification of pruriceptors [11,23]. Tlx3 transcripts are suppressed across all developmental timepoints analyzed when compared to wild type (Figure 7C). To further evaluate the likelihood that Tlx3 dysregulation could affect pruriceptor gene expression in Tmem184b mutants, we asked whether loss of Tlx3 and loss of Tmem184b caused similar expression profiles. Strikingly, more than half of the differentially expressed genes (adjusted p<0.01) from Tmem184b adult DRG were also disrupted by Tlx3 conditional loss in nociceptors (Fig. 7D and Table, Supplementary Digital Content 4) [11]. This analysis identifies Tlx3 as a possible link between early and late events regulated by Tmem184b.

Our data suggest a plausible model for the effects of Tmem184b on pruriception (Fig. 7E). In this model, Tmem184b first acts early in development to ensure proper expression of Wnt signaling pathway components. Following activation of Wnt signaling, both early-acting (Neurogenins) and later-acting (Tlx3) transcriptional programs are launched. Finally, extrinsic signals coordinate with Tlx3 in late embryonic and early postnatal development to execute proper DRG subtype gene expression. Tmem184b may also play a role later in pruriceptive maintenance and in proper receptor recycling at sensory terminals.

## Discussion

Our data identify a significant role for *Tmem184b* in the control of pruriception. *Tmem184b-mutant* DRGs show reduction of pruriceptor transcripts in the NP3/C2 population. Of note, mRNA expression of the IL-31 receptor subunit, *Il31ra*, and its co-receptor, *Osmr*, are both strongly decreased in the absence of *Tmem184b*. We see reduced IL-31-induced scratching as well as reduced calcium entry in individual neurons in response to IL-31 application in mutant mice. IL-31-mediated itch is central to the development of atopic dermatitis as well as allergic asthma, and thus the expression and function of its receptor is of significant medical interest [6,16,19]. Our data identify a novel mechanism promoting the expression of the IL-31 receptor on sensory neurons and ultimately impacting IL-31 induced behaviors.

While we initially hypothesized that the adult RNAseq and behavior results were caused by a loss of NP3/C2 neurons, further calcium imaging analysis suggests this may not be the case. IL31-responsive neurons still exist and still express some IL31RA (at least within the cell body), but the IL-31 evoked rise in intracellular calcium is reduced. Taken together with prior work suggesting that Tmem184b can be found on recycling endosomes within sensory neurons, we can speculate that it may have a role in maintenance or recycling of the IL31RA/OSMR heterodimeric receptor at terminals. This possibility should be examined in the future when better reagents are available to detect Tmem184b in tissues.

Our data from embryonic neurons indicate that *Tmem184b* promotes the expression of Wnt signaling components. At E13, Tmem184b mutants show substantial reductions in transcripts encoding critical developmental factors including *Wnt3a, Neurog1*, and *Neurog2*. Wnt signaling is critical for early neural crest development, and in stem cells, Wnt signaling induces *Neurog1, Neurog2*, and *Pou4f1* (*Brn3a*) expression [8,14,18]. These factors are essential for the establishment and differentiation of neurons within sensory ganglia [17]. It is perhaps surprising that sensory ganglia even form at all under these conditions; however, Tmem184b mutant mice form normal cohorts of nociceptors and pruriceptors. It could be that even low levels of these transcripts, alongside other developmental mechanisms, allow the full cohort of neurons to emerge but do not fully support normal adult subtype gene expression.

Based on this evidence, we believe that *Tmem184b* acts early in sensory development to promote the proper adult function of nociceptive and pruriceptive neurons. One possibility is that TMEM184B activity, through Wnt signaling, promotes the subsequent activation of the homeodomain transcription factor, *Tlx3. Tlx3* expression is reduced throughout our developmental timecourse in *Tmem184b* mutant mice compared to wild type. Further supporting its involvement, *Tlx3* nociceptor-specific knockouts show loss of pruriceptive populations. We show that pruriceptive markers showing reduced expression in *Tlx3*-mutant DRGs significantly overlap with *Tmem184b*-affected genes [11,23]. Intriguingly, the effect of blockade of Wnt secretion on the morphology of the mouse neuromuscular junction is strikingly similar to that seen in *Tmem184b-mutant* mice, further implicating a link between *Tmem184b* and Wnt signaling [2,29]. Future experiments should test how TMEM184B activity and appropriate Wnt signaling in early sensory development are associated.

We do not yet know the biological activity of TMEM184B, although its membership in the Transporter-Opsin-GPCR (TOG) superfamily of proteins [33] suggests a few exciting possibilities. Given all available evidence, including the presence of *Tmem184b* on recycling endosomes in adult sensory neurons [2], we suspect that *Tmem184b* may be involved in the availability of receptors on the cell surface, which would translate into reduced signaling and transcriptional output by pathways downstream of these receptors. For example, it is possible that TMEM184B activity influences both transcript expression and surface presentation of Wnt receptors during embryogenesis. Identifying the signaling pathways and cell biological processes activated by TMEM184B as neural crest cells take on neural identities and generate final fates should be a high priority for future studies. Because TMEM184B is known to contribute to synapse maintenance and axon degeneration following injury, the transcriptional dysregulation identified in our work may have important ramifications in maintaining neural connectivity.

In summary, we show that *Tmem184b* activity critically affects the function of pruriceptive neurons in mouse DRG and that this effect is likely due to its ability to induce components of Wnt signaling during neurogenesis. Our data illuminates a new key regulatory step in the processes controlling the establishment of diversity in the somatosensory system.

## Author Contributions

E.L. and M.B. designed the experiments. E.L. and M.B. performed RNAseq analysis and calcium imaging and analysis. T.C., J.F., E.L., C.M. and M.B. performed behavioral experiments. E.L., M.M., T.C, B.M., N.K., E.W., C.J., H.G. and M.B. analyzed behavior videos. T.L. consulted on experimental design and execution. H.H. and R.K. supported experiments in their laboratories and offered guidance on the project. E.G.L. and M.R.C.B. wrote the manuscript.

## Acknowledgements and Conflicts of Interests

We would like to thank Dr. Aubin Moutal for assistance with adult DRG cultures and calcium imaging. We would also like to thank Kai Lindstedt, Shanshan Zhang, Nathan Aviles, and Dr. Dean Billheimer for statistical consultation. Dr. Bhattacharya reports grants from NIH (R01NS105680) and the Muscular Dystrophy Association supporting the conduction of this study. This work was also supported by the University of Arizona (Technology and Research Incentive Fund), In addition, M.R.C.B. and T.M.L-M. have a patent UNIA 20.01 PCT pending. Beyond these, we have no conflicts of interest.

**Table, Supplemental Digital Content 1.** Table showing results of DESeq differential gene expression analysis between adult DRG of wild type and Tmem184b gene trap mice (n=4 per genotype). Genes for which adjusted p values are less than 0.05 are shown.

**Table, Supplementary Digital Content 2. Complete analysis of cellular nociceptive and pruriceptive responses by calcium imaging.**

Agonists concentrations were 1μM IL-31, 200μM chloroquine, 2.5μM LY344864, 250μM CYM5442, 5mM β-alanine, 200μM AITC, and 1μM capsaicin. Receptors for each agonist are given in parentheses. To estimate means of the proportions of neurons responding to each agonist, weighted linear regression was performed. Data from this analysis includes percent responders (weighted mean), standard error, and the regression p-values across genotypes. Weighted linear regression was also performed on each agonist group to estimate the mean peak response of ratiometric fluorescence above baseline and to calculate p-values. Peak analysis was limited to neurons imaged with ratiometric imaging (Fura-2 dye) for all agonists.

**Table, Supplementary Digital Content 3.** Genes and pathways of known developmental or neuronal importance and their expression (normalized counts as reported by DESeq2) at E13, P0, and P10. Highlights indicate genes for which at least one timepoint shows a significant difference between genotypes.

**Table, Supplementary Digital Content 4.** List of differentially expressed genes in Tlx3F/F, NaV1.8-Cre mice, analyzed using our pipeline and used for comparison to Tmem184b mutant ganglia in Figure 7.

