## Supplementary material for "TMEM184b is necessary for IL-31 induced itch": Supp Table 2

| Agonist (receptor) | Genotype | Number of Mice | Number of Responders | Number of Neurons | Percent Responders (Weighted Mean) | Std. Error | Percent Responders p-value |
| --- | --- | --- | --- | --- | --- | --- | --- |
| Chloroquine (MRGPRA3) | WT | 3 | 53 | 757 | 7.001 | 3.951 | 0.130 |
|  | Mut | 3 | 109 | 751 | 14.51 | 2.799 |  |
| LY344864 (HTR1F) | WT | 5 | 47 | 1240 | 3.79 | 1.062 | 0.208 |
|  | Mut | 5 | 28 | 1243 | 2.253 | 0.764 |  |
| CYM5442 (S1PR1) | WT | 4 | 57 | 967 | 5.894 | 1.146 | 0.059 |
|  | Mut | 4 | 34 | 1053 | 3.229 | 0.793 |  |
| β-alanine (MRGPRD) | WT | 3 | 169 | 1049 | 16.11 | 4.506 | 0.992 |
|  | Mut | 4 | 212 | 1312 | 16.16 | 3.003 |  |
| AITC (TRPA1) | WT | 5 | 304 | 1140 | 26.67 | 12.62 | 0.991 |
|  | Mut | 4 | 329 | 1241 | 26.51 | 8.73 |  |
| Capsaicin (TRPV1) | WT | 4 | 662 | 1105 | 59.91 | 6.378 | 0.128 |
|  | Mut | 4 | 594 | 1221 | 48.65 | 4.396 |  |

**Table 1. Complete analysis of cellular nociceptive and pruriceptive responses by calcium imaging.**

Agonists concentrations were 200μM chloroquine, 2.5μM LY344864, 250μM CYM5442, 5mM β-alanine, 200μM AITC, and 1μM capsaicin. Receptors for each agonist are given in parentheses. To estimate means of the proportions of neurons responding to each agonist, weighted linear regression was performed. Data from this analysis includes percent responders (weighted mean), standard error of the mean estimate, and the regression p-values across genotypes.
